# A flexible, efficient binomial mixed model for identifying differential DNA methylation in bisulfite sequencing data

**DOI:** 10.1101/019562

**Authors:** AJ Lea, J Tung, X Zhou

**Author notes:** These authors contributed equally to this work (J Tung, X Zhou). Other author.

## Abstract

Identifying sources of variation in DNA methylation levels is important for understanding gene regulation. Recently, bisulfite sequencing has become a popular tool for investigating DNA methylation levels. However, modeling bisulfite sequencing data is complicated by dramatic variation in coverage across sites and individual samples, and because of the computational challenges of controlling for genetic covariance in count data. To address these challenges, we present a binomial mixed model and an efficient, sampling-based algorithm (MACAU: Mixed model association for count data via data augmentation) for approximate parameter estimation and *p*-value computation. This framework allows us to simultaneously account for both the over-dispersed, count-based nature of bisulfite sequencing data, as well as genetic relatedness among individuals. Using simulations and two real data sets (whole genome bisulfite sequencing (WGBS) data from *Arabidopsis thaliana* and reduced representation bisulfite sequencing (RRBS) data from baboons), we show that our method provides well-calibrated test statistics in the presence of population structure. Further, it improves power to detect differentially methylated sites: in the RRBS data set, MACAU detected 1.6-fold more age-associated CpG sites than a beta-binomial model (the next best approach). Changes in these sites are consistent with known age-related shifts in DNA methylation levels, and are enriched near genes that are differentially expressed with age in the same population. Taken together, our results indicate that MACAU is an efficient, effective tool for analyzing bisulfite sequencing data, with particular salience to analyses of structured populations. MACAU is freely available at www.xzlab.org/software.html.

**Author Summary:** DNA methylation is an important epigenetic modification involved in regulating gene expression. It can be measured at base-pair resolution, on a genome-wide scale, by coupling sodium bisulfite conversion with high-throughput sequencing (a technique known as ‘bisulfite sequencing’). However, the data generated by such methods present several challenges for statistical analysis. In particular, while the raw data generated from bisulfite sequencing experiments are read counts, they are often converted to proportions for ease of modeling, resulting in loss of information. Furthermore, although DNA methylation levels are known to be heritable—and are thus affected by kinship and population structure—existing approaches for modeling bisulfite sequencing data fail to account for this covariance. Such failure can lead to spurious associations and reduced power. Here, we present a new approach that models bisulfite sequencing data using raw read counts, while also taking into account population structure and other sources of data over-dispersion. Using simulations and two real data sets (publicly available data from *Arabidopsis thaliana* and newly generated data from *Papio cynocephalus),* we demonstrate that our model provides well-calibrated *p*-values and improves power compared with previous methods. In addition, the DNA methylation patterns identified by our method agree with those reported in previous studies.

## Introduction

DNA methylation — the covalent addition of methyl groups to cytosine bases — is a major epigenetic gene regulatory mechanism observed in a wide variety of species. DNA methylation influences genome-wide gene expression patterns, is involved in genomic imprinting and X-inactivation, and functions to suppress the activity of transposable elements [1–3]. In addition, DNA methylation is essential for normal development. For example, mutant *Arabidopsis* plants with reduced levels of DNA methylation display a range of abnormalities including reduced overall size, altered leaf size and shape, and reduced fertility [4–6]. In humans, DNA methylation levels are strongly linked to disease, including major public health burdens such as diabetes [7,8], Alzheimer’s disease [9,10], and many forms of cancer [7,11–15]. Together, these observations point to a central role for DNA methylation in shaping genome architecture, influencing development, and driving trait variation. Consequently, there is substantial interest in identifying the genetic [16–19] and environmental [20–23] factors that shape DNA methylation levels. Progress toward this goal requires statistical approaches that can handle the complexities of real world, population-based datasets. Here, we present one such approach, designed specifically for analyses of differential methylation levels in bisulfite sequencing datasets.

High-throughput bisulfite sequencing approaches, which include whole genome bisulfite sequencing (WGBS or BS-seq) [24], reduced representation bisulfite sequencing (RRBS) [25,26], and sequence capture followed by bisulfite conversion [27,28], are used to estimate genome-wide DNA methylation levels at base-pair resolution. All such methods rely on the differential sensitivity of methylated versus unmethylated cytosines to the chemical sodium bisulfite. Specifically, sodium bisulfite converts unmethylated cytosines to uracil (and ultimately thymine following PCR), while methylated cytosines are protected from conversion. Estimates of DNA methylation levels for each cytosine base can thus be obtained directly from high-throughput sequencing data by comparing the number of C’s (reflecting an originally methylated version of the base) versus T’s (reflecting an originally unmethylated version of the base) at that position in the mapped reads.

The raw data produced by bisulfite sequencing methods are therefore count data, in which both the number of methylated reads and the total coverage at a site contain useful information. Higher total coverage corresponds to a more reliable estimate of the true DNA methylation level, which, in a typical experiment, can vary dramatically across individuals and sites (e.g., by several orders of magnitude: S1 Figure). Many commonly used methods for testing for differential methylation (whether by genotype, environmental predictor, or experimental perturbation) ignore this variability by converting counts to percentages or proportions (e.g., t-tests, Mann-Whitney U tests, linear models, and all tools initially designed for array-based data [29,30]; Table 1). Thus, a site at which 5 of 10 reads are designated as methylated (i.e., read as a cytosine) is treated identically to a site at which 50 of 100 reads are designated as methylated. This assumption reduces the power to uncover true predictors of variation in DNA methylation levels, because it treats noisy measurements the same way as accurate ones.

**Table 1.** Approaches for identifying differentially methylated loci in bisulfite sequencing data sets.

| Statistical method | Directly models counts? | Controls for biological covariates? | Controls for genetic covariance? | Programs that implement the method |
| --- | --- | --- | --- | --- |
| t-test or Wilcoxon rank-sum test | No | No | No | R and many others |
| Fisher's exact test | Yes | No | No | R and many others |
| Binomial regression | Yes | Yes | No | R and many others |
| Linear regression | No | Yes | No | R and many others |
| Beta-binomial model | Yes | Some<sup>1</sup> | No | DSS [31], MOABS [32], RadMeth [33] |
| Linear mixed model | No | Yes | Yes | GEMMA [34], EMMA [35], EMMAX [36], FaST-LMM [37] |
| Binomial mixed model | Yes | Yes | Yes | MACAU |

<sup>1</sup>Only RadMeth; the implementations of the beta-binomial model in MOABS and DSS do not allow the user to control for covariates.

To address this problem, several recently introduced methods for differential DNA methylation analysis implement a beta-binomial model (e.g., ‘DSS: Dispersion Shrinkage for Sequencing data’ [31], ‘RADMeth: Regression Analysis of Differential Methylation’ [33], and ‘MOABS: Model Based Analysis of Bisulfite Sequencing data’ [32]). These methods model the binomial nature of bisulfite sequencing data, while taking into account the well-known problem of over-dispersion in sequencing reads. Because these methods work directly on count data, they can reliably account for variation in read coverage across sites and individuals. Consequently, beta-binomial methods consistently provide increased power to detect true associations between genetic or environmental sources of variance and DNA methylation levels [31–33].

However, methods based on beta-binomial models only account for over-dispersion due to independent variation, making them unsuited for data sets containing population structure or related individuals. Accounting for genetic relatedness is important because genetic variation can exert strong and pervasive effects on DNA methylation levels [17,19,38,39]. In humans, methylation levels at more than ten thousand CpG sites are influenced by local genetic variation [18], and DNA methylation levels in whole blood are 18%-20% heritable on average, with the heritability estimates for the most heritable loci (top 10%) averaging around 68% [38,39]. As a result, DNA methylation levels will frequently covary with kinship or population structure, and failure to account for this covariance could lead to spurious associations or reduced power to detect true effects. This phenomenon has been extensively documented for genotype-phenotype association studies [35,36,40–42], and controlling for genetic covariance between samples is now a basic requirement for genome-wide association studies. Similar logic applies to analyses of gene regulatory phenotypes and studies of gene expression variation often do take genetic structure into account by using mixed model approaches [43–45]. However, despite growing interest in environmental epigenetics and epigenome-wide association studies (EWAS), none of the currently available count-based methods appropriately control for genetic effects on DNA methylation levels in bisulfite sequencing data (Table 1). Consequently, even though count-based methods have been shown to be more powerful, recent bisulfite sequencing studies have turned to linear mixed models to deal with the confounding effects of population structure [19,46].

To address this gap, we present a binomial mixed model (BMM) for identifying differentially methylated sites that directly models raw read counts while accounting for both covariance between samples and extra over-dispersion caused by independent noise. We also present an efficient, sampling-based inference algorithm to accompany this model, called MACAU (Mixed model association for count data via data augmentation). MACAU works directly on binomially distributed count data from any high-throughput bisulfite sequencing method (e.g., WGBS, RRBS, targeted sequence capture) and uses random effects to not only model over-dispersion (as in the standard beta-binomial approach [47]), but also to model relatedness/population structure. Hence, MACAU enables users to identify differentially methylated sites in a wide variety of settings, with little cost to power even when genetic effects on DNA methylation levels are negligible.

We compared MACAU’s performance with currently available methods under two realistic scenarios, using both real bisulfite sequencing data sets (WGBS and RRBS) and simulations parameterized based on properties of real data. In the first scenario, we analyzed publicly available data from *Arabidopsis thaliana* [48] to show that, when a predictor variable of interest is correlated with population structure, MACAU provides better control of type I error than existing methods. This setting is particularly relevant to understanding geographic variation in DNA methylation levels (e.g., [19,48–50]) and for identifying genetic or environmental predictors of DNA methylation in structured samples (e.g., [50,51]). In the second scenario, we used newly generated RRBS data from wild baboons *(Papio cynocephalus)* to demonstrate that MACAU also provides increased power to detect truly differentially methylated sites in the presence of kinship—a condition that often holds in analyses of natural populations (e.g., [48,52,53]) and in tests for epigenetic discordance between siblings [22,53–55]. As interest in epigenome-wide association studies (EWAS), environmental epigenetics, and the epigenetic correlates of disease grows, these types of complex data sets will become increasingly common.

## Results

### The binomial mixed model and the MACAU algorithm

Here, we briefly describe the model and the algorithm. Additional information is provided in the Supplementary Information Text File, which includes details on the model, inference method, and algorithm (including descriptions of the data augmentation approach and efficient MCMC sampling steps).

To detect differentially methylated sites, we model each potential target of DNA methylation individually (i.e., we model each CpG site one at a time) as a function of x, a predictor variable of interest. Here, x could be a genotype value, as in methylation QTL mapping analyses; an environmental predictor of interest, such as temperature, chemical exposure, or social environment; an individual characteristic, such as age or sex; or an experimental perturbation, as in a treatment-control design. For each site, we consider the following binomial mixed model (BMM): 

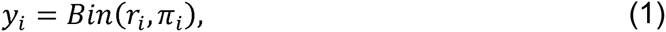
where *r_i_* is the total read count for *i*th individual; *y_i_* is the methylated read count for that individual, constrained to be an integer value less than or equal to *r_i_* and *π_i_* is an unknown parameter that represents the underlying proportion of methylated reads for the individual at the site. We use a logit link to model as a linear function of several parameters: 

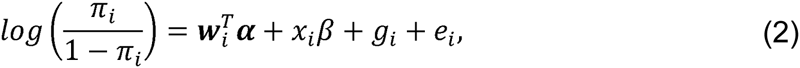

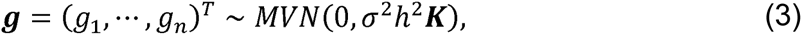

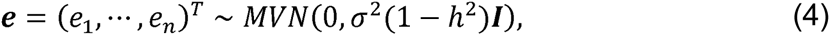
where, for a data set including *c* covariates and *n* individuals, ***w****_i_* is a *c*-vector of covariates including an intercept; *α* is a *c*-vector of corresponding coefficients; *x_i_* is the predictor of interest for individual *i* and *β* is its coefficient; ***g*** is an *n*-vector of genetic random effects that model correlation due to population structure or kinship; MVN denotes the multivariate normal distribution; ***e*** is an *n*-vector of environmental residual errors that model independent variation; ***K*** is a known *n* by *n* relatedness matrix that can be calculated based on pedigree or genotype data; ***I*** is an *n* by *n* identity matrix; *σ*^2^*h*^2^ is the genetic variance component; *σ*^2^(1 − *h*^2^) is the environmental variance component; and *h*^2^ is the heritability of the logit transformed methylation proportion (i.e. *logit*(***π***)). Note that ***K*** has been standardized to ensure *tr(****K****)/n* = 1, so that *h*^2^ lies between 0 and 1 and can be interpreted as heritability (see [56]; *tr* denotes the trace norm).

Both ***g*** and ***e*** model over-dispersion (i.e., the increased variance in the data that is not explained by the binomial model). However, they model different aspects of over-dispersion: ***e*** models the variation that is due to independent environmental noise (a known problem in data sets based on sequencing reads [57–60], including analyses of read proportions [61]), while ***g*** models the variation that is explained by kinship or population structure. Effectively, our model improves and generalizes the beta-binomial model by introducing this extra ***g*** term to model individual relatedness due to population structure or stratification. In the absence of ***g***, our model becomes similar to other beta-binomial models previously developed for modeling count data [31,33,47,62].

We are interested in testing the null hypothesis that the predictor of interest has no effect on DNA methylation levels: ***H*_0_**: *β* = 0. This test requires obtaining the maximum likelihood estimate 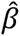 from the model. Unlike its linear counterpart, estimating 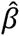 from the binomial mixed model is notoriously difficult, as the joint likelihood consists of an *n*-dimensional integral that cannot be solved analytically [63,64]. Standard approaches rely on numerical integration [65] or Laplace approximation [66,67], but neither strategy scales well with the increasing dimension of the integral, which in our case is equal to the sample size. Because of this problem, standard implementations of binomial mixed models often produce biased estimates and overly narrow (i.e., anti-conservative) confidence intervals [68–72]. To overcome this problem, we instead use a Markov chain Monte Carlo (MCMC) algorithm-based approach for inference, using un-informative priors for the hyperparameters *h*^2^ and *σ*^2^. After drawing accurate posterior samples of *β*, we rely on the asymptotic normality of both the likelihood and the posterior distributions [73] to obtain the approximate maximum likelihood estimate 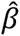 and its standard error se(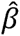). This procedure allows us to construct approximate Wald test statistics and *p*-values for hypothesis testing. Despite the stochastic nature of the procedure, the MCMC errors are small enough to ensure stable *p*-value computation across multiple MCMC runs (S2 Figure). We note that with reasonably large sample sizes (n=50 or more), the resulting p-values are also robust to prior perturbation on hyper-parameters (S3 Figure); however, all results reported here are based on calculations with un-informative priors.

In addition to the approximate inference procedure described above, we also developed a novel MCMC algorithm based on an auxiliary variable representation of the binomial distribution for efficient, approximate *p*-value computation [74–76] (see SI Text File Section 2: Inference Method Overview and SI Text File Section 3.1: Data Augmentation for more details). We did so to reduce the heavy computational burden of standard MCMC algorithms, which would otherwise be prohibitive in terms of run time for large datasets. Building on the auxiliary variable representation, our main technical contribution is a new framework that approximates the distribution of the auxiliary variables (S4 Figure, S1-S2 Tables) while simultaneously taking advantage of recent innovations for fitting mixed effects models [34,35,37,77] (see SI Text File Sections 3.2 and 3.3). This framework reduces per-MCMC iteration computational complexity from cubic to quadratic with respect to the sample size, and results in an approximate *n*-fold speed up in practice compared with the popular Bayesian software MCMCglmm [78], where *n* is the sample size (S5 Figure, S3 Table; we note that this speed-up is generalizable to other GLMM problems as well). Our implementation of the BMM is therefore efficient for data sets ranging up to hundreds of samples and millions of sites, as computational complexity scales only linearly with respect to the number of analyzed sites (S5 Figure).

Because our model effectively includes the beta-binomial model as a special case, we expect it to perform similarly to the beta-binomial model in settings in which population structure is absent (we say “effectively” because the beta-binomial model uses a beta distribution to model independent noise while we use a log-normal distribution). However, we expect our model to outperform the beta binomial in settings in which population structure is present. In addition, in the presence of population stratification, we expect the beta-binomial model to produce inflated test statistics (thus increasing the false positive rate) while our model should provide calibrated ones. Below, we test these predictions using two different bisulfite sequencing data sets. We begin with simulations in which the true value of *β* is known, and the over-dispersion parameter and genetic covariance between samples are motivated by the real data sets. We also motivate our choice of simulated sample sizes based on real bisulfite sequencing data sets, which currently range from ~20 − 150 samples [19,26,46,53,79–82]. However, because sample sizes are only likely to grow in the future, for the data set types of most direct interest (i.e., those that contain population structure and heritable DNA methylation levels) we further consider sample sizes that are much larger than currently represented in the literature (n = 500 and n = 1000). Finally, we apply our model directly to the real data.

### Count-based models perform well in the absence of genetic effects on DNA methylation levels

We first compared the performance of the BMM implemented in MACAU with the performance of other currently available methods for analyzing bisulfite sequencing data in the absence of genetic effects. Intuitively, we expected MACAU and the beta-binomial model to perform similarly, and we expected both methods to outperform those that first transform the raw count data. To test our prediction, we simulated the effect of a predictor variable on DNA methylation levels across 5000 CpG sites (4500 true negatives and 500 true positives). Motivated by our analysis of age effects on DNA methylation levels in the baboon RRBS data set (below), we conducted this simulation by sampling from a distribution of known age values from the same baboon population. For all simulations, we set the effect of genetic variation on DNA methylation levels equal to zero, which is equivalent to setting either (i) the heritability of DNA methylation levels to zero (unlikely based on prior findings [38,39]), or (ii) studying completely unrelated individuals in the absence of population structure. To explore MACAU’s performance across a range of conditions, we simulated age effects on DNA methylation levels across three effect sizes (percent of variance in DNA methylation explained (PVE) = 5%, 10%, or 15%) and three sample sizes (n = 20, 50, and 80). These values capture the majority of effect sizes and sample sizes documented in recent genome-wide bisulfite sequencing studies (e.g., [45,52,53,83]).

Because age is naturally modeled as a continuous variable, we focused our comparisons only on approaches that could accommodate continuous predictor variables (comparisons in which we artificially binarized age, which allowed us to include a larger set of approaches, are shown in S6 Figure and S7 Figure for cases excluding and including genetic effects on DNA methylation, respectively; however, binarizing a truly continuous variable consistently results in poorer performance: see S6 Figure versus S9 Figure). Specifically, in addition to the BMM implemented in MACAU, we considered the performance of a beta-binomial model, a binomial model, a linear model, and a linear mixed model (implemented in the software GEMMA [34]). For the linear and linear mixed model case, methylation proportions were quantile normalized to a standard normal prior to modeling (see Methods and S8 Figure for parallel results using logit, M-value, and arcsin(sqrt) transformations prior to linear mixed modeling as alternatives to quantile normalization). As expected, we found that MACAU performed similarly to the beta-binomial model, and that these two approaches consistently detected more true positive age effects on DNA methylation levels (at a 10% empirical FDR) than all other methods (S9 Figure). For example, in the “easiest” case we simulated (PVE = 15%, n = 80), we found that the beta-binomial model detected 30% of simulated true positives, while the BMM implemented in MACAU detected 27.8%. The slight loss of power in the BMM is a consequence of the smaller degrees of freedom caused by the additional genetic variance component. In comparison, the linear model detected 21.2% of true positives; the linear mixed effects model, 14%; and the binomial model, 8.4% (S9 Figure). Although it is often used to test for differential methylation [53,84,85], the binomial model exhibits low power when an empirical FDR is used to control for multiple hypothesis testing due to poor type I error calibration, as has been previously reported [33]. Area under a receiver operating characteristic curve (AUC) was also consistently very similar between the beta-binomial and MACAU (S9 Figure), although the advantage of the count-based methods was less clear by this measure. This reduced contrast is because AUC is based on true positive-false positive trade-offs across the entire range of *p*-value thresholds: methods can consequently yield high AUCs even when they harbor little power to detect true positives at FDR thresholds that are frequently used in practice. Taken together, our simulations suggest a general advantage to count-based models for samples that contain no genetic structure. Further, the differences in performance between the beta-binomial model and the BMM implemented in MACAU were consistently small in this setting (S9 Figure).

### Binomial mixed models control for false positive associations that arise from population structure: simulations and a real data example from *Arabidopsis*

We next evaluated each model’s performance in a more realistic setting, in which genetic covariance between samples could potentially confound tests for environmental or genetic effects on DNA methylation levels. As a case study example, we drew from publicly available phenotype data and SNP genotype data for 24 *Arabidopsis thaliana* accessions [86,87] in which leaf tissue samples had been recently subjected to whole genome bisulfite sequencing [48]. Among these accessions, a secondary dormancy phenotype (measured as the slope of the relationship between length of cold treatment and seed germination percentages [88]) is correlated with population structure (R^2^ = 0.38 against the first principal component of the genotype matrix for these accessions; p = 7.84 × 10^−4^; S10 Figure). Because secondary dormancy is associated with environmental conditions that are experienced after the seed has already dispersed, we have no expectation that secondary dormancy should be associated with DNA methylation levels in leaf tissue. Consequently, this data set provided the opportunity to evaluate calibration of Type I error (false positives) using MACAU, which controls for population structure, versus other available approaches.

To do so, we first used the true distribution of secondary dormancy characteristics and the true genetic structure among these 24 accessions to simulate a dataset that consisted entirely of null associations. Specifically, we simulated data sets (containing 4000 sites each) in which the secondary dormancy had no effect on DNA methylation levels, but the effect of genetic variation on DNA methylation levels was either moderate (h^2^ = 0.3) or large (h^2^ = 0.6). Thus, in these data sets, population structure could confound the relationship between the predictor variable (the capacity for secondary dormancy) and DNA methylation levels if not taken into account.

As predicted, we found that the BMM implemented in MACAU appropriately controlled for genetic effects on DNA methylation levels: whether DNA methylation levels were moderately (h^2^ = 0.3) or strongly (h^2^ = 0.6) heritable, MACAU did not detect any sites associated with secondary dormancy at a relatively liberal false discovery rate threshold of 20% (whether calculated against empirical permutations or calculated using the R package *qvalue* [32]). In addition, the *p*-value distributions for secondary dormancy effects on DNA methylation levels, in both simulations, did not differ from the expected uniform distribution (Fig. 1; Kolmogorov-Smirnov (KS) test when h^2^ = 0.3: D = 0.015, p = 0.909; when h^2^ = 0.6: D = 0.016, p = 0.874; genomic control factors: 0.90 when h^2^ = 0.3, 0.93 when h^2^ = 0.6). In contrast, when we analyzed the same simulated data sets with a beta-binomial model, we erroneously detected 2 CpG sites associated with secondary dormancy when heritability was set to 0.3, and 4 CpG sites when heritability was set to 0.6 (at a 20% FDR in both cases). More concerningly, the distributions of *p*-values produced by the beta-binomial model were significantly different from the expected uniform distribution and skewed towards low (significant) values (KS test when h^2^ = 0.3: D = 0.084, p = 1.75 × 10^−8^; when h^2^ = 0.6: D = 0.096, p = 2.80 × 10^−11^; genomic control factors: 1.18 when h^2^ = 0.3, 1.32 when h^2^ = 0.6). These results suggest an increasing problem with false positives as the heritability of DNA methylation levels increases (see S11 Figure for similar results when comparing a linear model to a linear mixed model).

**Fig. 1.** MACAU appropriately controls for genetic covariance in simulated and real WGBS data and eliminates false positive identification of differentially methylated sites. (A, B) The distribution of *p*-values for 4000 simulated true negative sites (n = 24 accessions; effect of secondary dormancy on DNA methylation levels = 0). For each simulation, h^2^ was set to 0.3 (A) or 0.6 (B). Simulated data were analyzed with a beta-binomial model or MACAU, and compared against the expected uniform distribution. (C) QQ-plots comparing the *p*-value distributions for (i) a model testing for effects of secondary dormancy on DNA methylation levels in real WGBS data, with quantiles plotted on the y-axis; and (ii) the same model when the secondary dormancy values were permuted across individuals, with quantiles plotted on the x-axis. The genomic control factor, λ, is shown for each set of results.

Notably, this problem should become more acute with increasing sample size, which provides greater power to detect false positives generated by this type of confounding [89]. Indeed, both increasing the simulated sample size and increasing the simulated correlation between the predictor variable and genetic structure produces increasingly poorly calibrated results. For example, when sample sizes were simulated from 25 up to 1000 individuals (and the heritability of DNA methylation levels was set to 0.6), we observed genomic inflation factors ranging from 1.03 – 3.49 for data sets analyzed with a beta-binomial (Fig. 2a). Not surprisingly, for a dataset of a fixed size, the beta-binomial genomic control factor increased as the confounding between population structure and the predictor variable of interest became more extreme (see S12a Figure for comparable results for a linear model). In contrast, when we analyzed the same simulated datasets with the BMM implemented in MACAU, the genomic control factors consistently ranged from 0.82 – 1.08, even when sample sizes were large and/or the correlation between population structure and the predictor variable was substantial (Fig. 2b; see S12b Figure for comparable results from a linear mixed model). Importantly, these differences in genomic control factors can translate into substantial differences in the results suggested by a given method. For example, when n = 1000 and the predictor variable is highly confounded with population structure (R^2^ = 0.5), a beta-binomial falsely identified 32% of sites in the data set as differentially methylated (10% FDR), while MACAU correctly identified no differentially methylated sites (10% FDR; S13 Figure).

**Fig. 2:** MACAU controls for genetic covariance in data sets that span a range of sample sizes and levels of correlation between population structure and a predictor variable of interest. Genomic control factor when simulated datasets (n=5000 sites per dataset; h^2^ = 0.6) were analyzed with either (A) a beta-binomial model or (B) a BMM implemented in MACAU. The correlation between the simulated predictor variable and the first principal component of genome-wide genotype data is plotted on the x-axis. Genotype data are for *Arabidopsis* accessions, as reported in [87].

To investigate the calibration of test statistics in the real data set, we then analyzed the relationship between the secondary dormancy phenotype and WGBS data for the 24 *Arabidopsis* accessions in which both phenotype and WGBS data were available (n = 830,676 CpG sites tested [32,33,34]). We again compared the performance of a simple linear model, a binomial model, a beta-binomial model, the BMM implemented in MACAU, and an LMM implemented in GEMMA. Further illustrating its poor handling of Type I error, the binomial model detected more than 100,000 secondary dormancy-associated sites at a 10% empirical FDR threshold, respectively, with a genomic control factor of 3.81. A beta-binomial model substantially improved over the binomial model, but still detected 39 secondary dormancy-associated sites at a 20% empirical FDR threshold, and 150 sites and 690 sites at a 10% or 20% FDR *qvalue* threshold, respectively (genomic control factor = 1.16). Given the clear confounding of population structure and secondary dormancy in this sample, as well as the results of our simulations, these associations are probably largely, if not completely, spurious. In contrast, MACAU, the linear mixed model (GEMMA), and the simple linear model did not identify any CpG sites associated with secondary dormancy, either at a 10% or a 20% false discovery rate threshold (Fig. 1 and S11 Figure; genomic control factors: MACAU – 0.89, GEMMA – 0.97, Linear model – 0.99). Based on our earlier simulations, the similarity of performance among the three approaches likely stems from different reasons: the linear model is poorly powered to detect positive hits with this sample size (either true positives or false positives); the linear mixed model controls for population structure but has low power to detect true associations; while MACAU combines both the increased power conferred by modeling the raw count data with appropriate controls for population structure (see Fig. 1 and results below).

### MACAU provides increased power to detect true positives in the presence of kinship: simulations based on data from baboons

In other data sets, a predictor variable of interest may not be confounded with genetic structure, but modeling genetic similarity between samples could reduce residual error variance and improve power. To investigate this scenario, we focused on the relationship between age and DNA methylation levels in a wild baboon population. Female baboons remain in their natal groups throughout their lives, producing relatedness values that are primarily due to matrilineal descent. The resulting genetic structure is one in which females tend to be more closely related to each other, on average, than males or male-female dyads [90], but in which not all females are related (because multiple matrilines co-reside in a single group). Data sets drawn from baboon populations therefore include a substantial number of unrelated individuals, but also some dyads that are genetically non-independent (i.e., relatives: S14 Figure).

To test the relative performance of different modeling approaches in this setting, we first simulated moderate to large genetic effects on DNA methylation levels (h^2^ = 0.3 and 0.6 respectively, as in the *Arabidopsis* simulation above) and relatedness values based on the observed distribution of relatedness values within baboon social groups (n = 80, 500, or 1000 baboons). We again simulated a range of non-zero effect sizes (percent variance explained by age = 5%, 10%, or 15%) for 500 true positive sites, and an effect size of zero for 4500 true negative sites.

In simulations in which age had a moderate effect on DNA methylation levels (PVE = 10%), MACAU detected 11.4% (when h^2^ = 0.3) and 20.6% (when h^2^ = 0.6) of simulated true positives at a 10% empirical FDR, and produced well calibrated *p*-values for sites with no simulated age effect (S15 Figure). In comparison, the beta-binomial model (the next best model) detected 8.2% and 10.4% of true positives, respectively (Fig. 3). As in the simulations, we again observed that a simple binomial model was prone to type I error, which resulted in failure to detect true age-associated sites when empirical FDRs were calculated against permuted data. Our additional simulations at PVE = 5% or PVE = 15%, and n = 500 or n = 1000, confirmed MACAU’s advantage over other methods across a range of conditions (S16-S17 Figure). As expected, the magnitude of this advantage was positively correlated with the heritability of DNA methylation levels.

**Fig. 3.** MACAU exhibits increased power to detect differential methylation when DNA methylation levels are heritable. Receiver operating characteristic (ROC) curves and true positive rates at a 10% false discovery rate threshold for simulated age effects on DNA methylation levels at (A-C) simulated sites with moderately heritable DNA methylation levels (h^2^ = 0.3) and (D-F) simulated sites with highly heritable DNA methylation levels (h^2^ = 0.6). Panels B and E are enlarged versions of panels A and D, respectively. They focus on false positive rates below 0.1, because the performance of alternative methods at low false positive rates tends to be most important to researchers in practice; that is, it is unlikely to matter if method performance is identical when accepting a 50% false positive rate, which would yield very poor inferential power. Each simulated dataset contained n=80 individuals and 5000 simulated CpG sites, with 500 true positives and 4500 true negatives. Here, we show results where the simulated percent variance explained by age = 10%. A binomial model could not detect true positives at a false positive rate below 0.10 (when h^2^ = 0.3) or below 0.9 (when h^2^ = 0.6); the binomial is therefore removed from panel B, and only shown for large false positive rates in panel E.

### Age-associated DNA methylation levels in wild baboons

Finally, we analyzed the new baboon RRBS data set for differential methylation patterns by age (n = 50, age range = 1.76 - 18.01 years in our sample, S4 Table). Because age-related effects on DNA methylation levels are well described, this approach allowed us to not only evaluate MACAU’s ability to detect differentially methylated sites, but also to identify known age-related signatures in DNA methylation data [38,39,91–93]. This data set included 433,871 CpG sites, enriched for putatively functional regions of the genome (e.g., genes, gene promoters, CpG islands, as expected in RRBS data sets [25,26]: S18 Figure; see also S19 Figure and S4 Table for additional information on data quality, including bisulfite conversion rates, *MspI* digest efficiency, correlation with gene expression levels, and methylation level distributions by genomic regions). As in our simulations, we found that MACAU provided increased power to detect age effects in the presence of familial relatedness. We detected 1.6-fold more age-associated CpG sites at a 10% empirical FDR using MACAU compared to the results of a beta-binomial model, the next best approach (1.4-fold more sites at a 20% empirical FDR; Fig. 4 and S20 Figure). This advantage was consistently observed across all FDR thresholds we considered, except for relatively low (<7.5%) empirical FDR thresholds, when all of the methods were very low powered as a result of the modest sample size.

**Fig. 4.** Age-associated CpG sites identified by MACAU in the baboon RRBS data. (A) The number of age-associated CpG sites detected at a given empirical FDR. The binomial model cannot detect age-associated sites at a false discovery rate below 0.20 and is consequently removed from the panel. (B) For age-associated sites detected by MACAU (at a 10% FDR), the proportion of sites that gain or lose methylation with age is shown by genomic region. Positive = DNA methylation levels increase with age; Negative = DNA methylation levels decrease with age. (C) Age-associated CpG sites detected using MACAU (10% FDR) are more likely to fall near genes that are expressed in whole blood, compared to the background set of CpG sites near genes (**p < 10^−10^). Further, age-associated CpG sites are more likely to occur near genes that are differentially expressed (DE) with age, compared to CpG sites near genes that are not DE with age (*p = 0.032).

We performed several analyses to investigate the likely validity and functional importance of the age-associated CpG sites we identified. Based on the results of previous studies, we expected that age-associated sites in CpG islands would tend to gain methylation with age [92,93], while sites in other regions of the genome (e.g., CpG island shores, gene bodies) would tend to lose methylation with age [92,93]. In addition, we expected that, in whole blood, bivalent/poised promoters should gain DNA methylation with age, while enhancers should lose methylation with age (as discussed in [91,92,94]). Finally, we expected that stretches of differentially methylated sites (i.e., differentially methylated regions, or DMRs) would tend to occur in or near CpG islands and CpG shores, potentially altering how steeply methylation levels change between islands and their surrounding shelves (e.g., [95]).

Our results conformed to these patterns: sites in CpG islands tended to gain methylation with age (71.4% of sites were positively correlated with age); and sites in promoters, CpG island shores, and gene bodies tended to lose methylation with age (72.7%, 75.4%, and 75.2% of sites were negatively correlated with age, respectively; Fig. 4). In addition, we found that positively correlated, age-associated sites were highly enriched in chromatin states associated with bivalent/poised promoters (as defined by the Roadmap Epigenomics Project [96]). Specifically, age-associated CpG sites in bivalent/poised promoters were 3.4 times more likely to show increases in DNA methylation with age, compared to age-associated CpG sites in other regions (p < 10^−10^, Fisher’s exact test). Negatively correlated age-associated sites (i.e., sites where DNA methylation levels decreased with age) were strongly enriched in enhancers (defined as sites either marked by H3K4me1 in human PBMCs [97] or sites within chromatin states annotated as ‘enhancers’ by the Roadmap Epigenomics Project [96], p = 2 × 10^−4^, Fisher’s exact test). Finally, we detected 142 age-related DMRs, the majority of which were found in CpG islands, shores, and bridging islands and shores (S21 Figure and S5 Table).

We also reasoned that true positive age-associated CpG sites should contain information about age-associated gene expression levels. To test this hypothesis, we turned to previously generated whole blood RNA-seq data [43] from the same baboon population (n = 63; only four baboons in the RNA-seq data set were also included in the DNA methylation data set). Overall, we observed a strong enrichment of differentially methylated CpG sites in or near (within 10 kb) blood-expressed genes (n = 12,018 genes), compared to the background set of all CpG sites near genes (Fisher’s exact test, p < 10^−10^). Further, CpG sites near age-associated genes (n = 1396 genes, 10% FDR) were 30.5% more likely to be differentially methylated with age compared to the background set of all CpG sites near genes (Fisher’s exact test, p = 0.032; Figure 4). Notably, this enrichment was almost always stronger for the set of differentially methylated sites identified by MACAU than for the same number of top sites identified when running the linear model, linear mixed model, binomial, or beta-binomial approaches, across different FDR thresholds (S22 Figure).

## Discussion

DNA methylation levels can have potent effects on downstream gene regulation, and, in doing so, can shape key behavioral, physiological, and disease-related phenotypes [7,20,98–100]. These observations have motivated an increasing number of DNA methylation studies in humans and other organisms, highlighting the need for sophisticated statistical methods that can accommodate the complexities of a broad array of data sets [19,46]. Here, we demonstrate that the binomial mixed model implemented in our software MACAU can (i) effectively control for confounding relationships between genetic background and a predictor variable of interest and (ii) provide increased power to detect true sources of variance in DNA methylation levels in data sets that contain kinship or population structure. In addition, MACAU provides increased flexibility over current count-based methods that cannot accommodate biological replicates (e.g., Fisher’s exact test), continuous predictor variables (e.g., DSS, MOABS, RadMeth), or biological or technical covariates (e.g., MOABS, DSS; see also Table 1). Given the increasing interest in both the environmental [21,101,102] and genetic [16,17,19,103] architecture of DNA methylation levels, we believe MACAU will be a useful tool for generalizing epigenomic studies to a larger range of populations. MACAU is particularly well suited to data sets that contain related individuals or population structure; notably, several major population genomic resources contain structure of these kinds (e.g., the HapMap population samples [104], the Human Genome Diversity Panel [105], and the 1000 Genomes Project in humans [106]; the Hybrid Mouse Diversity Panel in mice [107]; and the 1001 Genomes Project in *Arabidopsis* [108]).

Indeed, our results suggest MACAU is a useful tool even in data sets that are less affected by genetic structure, or when the heritability of DNA methylation levels is unclear. Because the beta-binomial model is effectively incorporated as a special case, MACAU exhibits only a slight loss of power relative to a beta-binomial model without genetic random effects when h^2^ = 0, while conferring better power and better test statistic calibration when h^2^ > 0 (S9, S16-S17 Figures, Fig. 1). Previous studies in humans have shown that, while the heritability of DNA methylation levels varies across loci, an appreciable proportion of loci are either modestly (h^2^ ≥ 0.3: 21.06% of all CpG sites) or highly (h^2^ ≥ 0.6: 6.95% of all CpG sites) heritable [39,109]. Further, DNA methylation QTLs are widespread across the genome [18,38,103]. Thus, because investigators will rarely have *a priori* knowledge of the heritability of DNA methylation levels at a given locus, and because the advantage of a beta-binomial model is small even when heritability is zero, we recommend applying MACAU in cases in which genetic effects on DNA methylation levels are poorly understood. In addition, our model provides a natural framework for incorporating the spatial dependency of DNA methylation levels across neighboring sites [110,111], which we expect to increase power even further [110,111]. However, we do note that, even with the efficient algorithm implemented here, fitting the binomial mixed model (or its extensions) remains more computationally expensive than other approaches for moderately sized datasets (S3 Table). While it remains appropriate for the sample sizes used in current studies (e.g., dozens to hundreds of individuals), or even larger with the support of a moderate-sized computing cluster (because MACAU is easily parallelizable with respect to sites), rapid increases in sample size—especially in the context of EWAS—strongly motivate additional algorithm development to scale up the binomial mixed model for data sets that include thousands or tens of thousands of individuals. This is particularly important given that methods tailored for other types of studies (e.g., quantile normalization followed by linear mixed modeling or *voom* + *limma,* both commonly used for RNA-seq) do not appear to translate well to bisulfite sequencing data sets (Figure S8; see Methods for additional information on the *voom + limma* comparison).

Although we developed MACAU with the analysis of bisulfite sequencing data in mind, we note that a count-based binomial mixed model may be an appropriate tool in other settings as well. For example, allele-specific gene expression (ASE) can be measured in RNA-seq data by comparing the number of reads originating from a given variant to the total number of mapped reads for that site [77,112–114]. Similarly, alternative isoform usage can be represented as a proportion of reads containing a non-constitutive exon versus the total reads for the same gene [47]. The structure of these data are highly similar to the structure of bisulfite sequencing data, which focus on counts of methylated versus total reads. Unsurprisingly, beta-binomial models have also emerged as one of the most popular methods for estimating both ASE values [114–116] and alternative isoform usage [47]. Researchers interested in the predictors of variation in either of these measures — which could include *trans-acting* genetic effects, environmental conditions, or properties of the individual (e.g., sex or disease status) — might also benefit from using MACAU. Recent work from the TwinsUK study motivates the need for such a model: Grundberg et al. demonstrated a strong heritable component to ASE levels [117], which could be effectively taken into account using the random effects approach implemented here.

Finally, linear mixed models have been recently proposed to account for cell type heterogeneity in epigenome-wide association studies focused on array data [118]. In this framework, the random effect covariance structure is based on overall covariance in DNA methylation levels between samples, which is assumed to be largely attributable to variation in tissue composition. MACAU provides a potential avenue for extending these ideas to sequencing-based data sets.

## Materials and Methods

### *Arabidopsis thaliana* whole genome bisulfite sequencing (WGBS) data set

We downloaded publicly available WGBS data generated by Schmitz et al. [48], as well as previously published SNP genotype data [87] and secondary dormancy data [86] for 24 *Arabidopsis* accessions. We used the SNP genotype data (specifically, 188,093 sites with minor allele frequency >5%) to construct a pairwise genetic relatedness matrix, K, as the product of a standardized genotype matrix *X*, or *K=XX^T^/p* [56], where genotypes were expressed as 0, 1, or 2 depending on the number of reference alleles for that site-sample combination. We used this estimate of *K* for both the simulations and our analyses of the real WGBS data.

In these analyses, we focused on CpG sites measured in ≥50% of accessions, and excluded sites that were constitutively hypermethylated (average DNA methylation level >0.90) or hypomethylated (average DNA methylation level <0.10, following [101,118]). We also excluded highly invariable sites (i.e., sites where the standard deviation of DNA methylation levels fell in the lowest 5% of the overall data set) and sites with very low coverage (i.e., sites where the mean coverage fell in the lowest quartile for the overall data set, below a mean of 3.34 reads). After filtering, the final data set consisted of 830,676 sites.

For the analysis of test statistic calibration as a function of sample size (Fig. 2), we also used *Arabidopsis* data, but simulated the phenotype data as a function of genetic covariance between the accessions. Genotype data were obtained from [87].

### Baboon reduced representation bisulfite sequencing (RRBS) data set Study subjects and sample collection

To investigate age effects on DNA methylation levels, in both real and simulated data sets, we drew on data and samples from a wild population of yellow baboons in the Amboseli ecosystem of southern Kenya. This population has been monitored for over four decades by the Amboseli Baboon Research Project (ABRP) [119], and the ages of animals born in the study population (n = 37; 74% of the data set) were therefore known to within a few days’ error. For animals that immigrated into the study population, ages were estimated from morphological features by trained observers (n = 13; 26% of the data set) [120]. Pairwise relatedness values were calculated based on previously collected microsatellite data (14 highly variable loci) [121,122], using the likelihood-based estimator of Lynch and Ritland [123] implemented in the program COANCESTRY [124]. Using the age and relatedness data sets, we simulated age effects on DNA methylation levels for either n = 20, 50, or 80 baboons from a single social group. For simulations with larger sample sizes, we extrapolated both age values and pairwise relatedness values from the n = 80 dataset to maintain the same level of age variation and genetic structure; notably, our results are highly stable in the face of realistic levels of noise in the estimate of *K* (S23 Figure). In addition, we used previously collected blood samples from the Amboseli population, paired with age and microsatellite genotype records, to investigate age effects on DNA methylation levels in a newly generated RRBS data set.

To generate the new RRBS data, we used whole blood samples collected from 50 animals (35 males and 15 females) by the ABRP between 1989 and 2011 following well-established procedures [43,125,126]. Briefly, animals were immobilized by an anesthetic-bearing dart delivered through a hand-held blow gun. They were then quickly transferred to a processing site for blood sample collection. Following sample collection, study subjects were allowed to regain consciousness in a covered holding cage until they were fully recovered from the effects of the anesthetic. Upon recovery, study subjects were released near their social group and closely monitored. Blood samples were stored at the field site or at an ABRP-affiliated lab at the University of Nairobi until they were transported to the United States.

Importantly, given the large range in sample collection dates, we observed no correlation between the age of our study subjects at sample collection and sample age (i.e., time since the collection date; Spearman rank correlation, p = 0.779). Further, to ensure that variation in sample collection dates did not influence our results, we also controlled for sample age as a covariate in our final analyses of the RRBS dataset (see *Analysis of age-related changes in DNA methylation levels).*

### RRBS data generation and low-level processing

Genomic DNA was extracted from whole blood samples using the DNeasy Blood and Tissue Kit (QIAGEN) according to the manufacturer’s instructions. RRBS libraries were created from 180 ng of genomic DNA per individual, following the protocol by Boyle et al. [25]. In addition, 1 ng of unmethylated lambda phage DNA (Sigma Aldrich) was incorporated into each library to assess the efficiency of the bisulfite conversion (>98% in all case: S4 Table). All RRBS libraries were sequenced using 100 bp single end sequencing on an Illumina HiSeq 2000 platform, yielding a mean of 28.97 ±8.97 million reads per analyzed sample (range: 9.59 – 79.78 million reads; Table S4).

We removed adaptor contamination and low-quality bases from all reads using the program TRIMMOMATIC [127]. We then mapped the trimmed reads to the olive baboon genome *(Panu* 2.0) using BSMAP, a tool designed for high-throughput DNA methylation data [128]. We used a Python script packaged with BSMAP to extract the number of reads as cytosine (reflecting an originally methylated base) and the total read count for each individual and CpG site. We performed the same set of filtering steps described for the *Arabidopsis* WGBS data set to produce our final data set for the baboons. Specifically, we excluded sites that were constitutively hypermethylated or hypomethylated, sites that were highly invariable, and sites that had low average coverage across individuals (in this case, the lowest quartile for mean coverage levels was 4.74 reads). The final filtered data set consisted of 433,871 CpG sites.

### Simulations

To simulate the methylated read counts and total read counts that result from WGBS and RRBS, we performed the following procedure:

First, we simulated the proportion of methylated reads for each site. To do so, we drew secondary dormancy values or age values, *x,* as the predictor of interest, from the actual values for the *Arabidopsis* accessions or from the baboon population, respectively. For simulations that focused on *Arabidopsis* data sets of various sizes (e.g., Figure 2), we simulated *x* and varied the degree to which it was confounded with population structure. Specifically, for each dataset (ranging from n=25 to n=1000 accessions) we performed principal components analysis on the SNP genotype data, and extracted the first principal component to capture the major axis of population structure (PC1). We then added environmental noise from a zero-centered normal distribution to achieve a correlation (R^2^) between the simulated phenotype and PC1 that reached the desired value (ranging from R^2^ = 0.1 to 0.5).

For each simulated data set, we simulated the DNA methylation level at each CpG site, *π,* as a linear function of *x* and its effect size, *β*. In addition, we included the effects of genetic variation *(g)* and random environmental variation (*e*), passed through a logit link (based on the model described in the Results section).

For the baboon RRBS and the *Arabidopsis* WGBS simulations, we determined *K* from 14 highly variable microsatellite loci or from the publicly available SNP data, as described above. For each simulation, we set *h*^2^ to 0, 0.3, or 0.6 to simulate non-heritable, modestly heritable, or highly heritable DNA methylation levels. We also estimated the variance term *σ*^2^ from the real data sets. Specifically, we took the mean estimate of *σ*^2^ across all sites (calculated in MACAU) for each real data set, and used this value as the fixed value of *σ*^2^ in the corresponding simulations.

Next, for each site, we simulated total read counts *r_i_* for each individual *i* from a negative binomial distribution that models the extra variation observed in the real data:

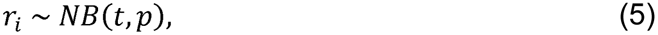

where *t* and *p* are site specific parameters estimated from the real data. Specifically, we generated 10,000 sets of *t* and *p* parameters by fitting a negative binomial distribution to the total read count data from 10,000 randomly selected CpG sites in the real baboon RRBS data set or the real *Arabidopsis* data set, using the function ‘fitdistr’ in the R package *MASS* [129]. To simulate counts for a given CpG site, we randomly selected one of these parameter sets to produce the total number of reads. Finally, we simulated the number of methylated reads for each individual at that locus *(y)* by drawing from a binomial distribution parameterized by the number of total reads (*r*) and the DNA methylation level *(π).*

### Comparison of MACAU to existing methods

For all simulated and real data sets, we used raw methylated and total read counts to compare the results of a beta-binomial model (using a custom R script), a binomial model (implemented via ‘glm’ in R), and the binomial mixed model implemented in MACAU. For computation time comparison, we used the MCMCglmm software, which also provides an implementation of a binomial mixed model [78]. In addition, we used the same count data to run a Fisher’s exact test (implemented in R), DSS [31], and RadMeth [33] in the subset of analyses that utilized these programs. To analyze simulated and real data sets using a linear model (implemented using *‘lm’* in R) or the linear mixed model implemented in GEMMA [34], we estimated DNA methylation levels by dividing the number of methylated reads by the total read count for each individual and CpG site. We then quantile normalized the resulting proportions for each CpG site to a standard normal distribution, and imputed any missing data using the K-nearest neighbors algorithm in the R package *impute* [130].

In addition to the quantile normalization approach, we also evaluated three other methods for transforming methylation proportions: a logit transformation, following [110]; the “M” value transformation (log_2_((methylated counts + *α*)/(unmethylated counts + *α*)), where *α* = 0.01, following [30]; and an arcsin square root transformation, following [131]. All four approaches produced qualitatively identical results (S8 Figure), so we elected to concentrate on the results from quantile normalization in the main text. Finally, we also tested the performance of a powerful, commonly used method for modeling RNA-seq data: the combination of the *voom* function for data weighting with *limma,* a linear model approach [132]. Our results indicated that *voom* + *limma* performs more poorly than even a simple linear model (S24 Figure), probably because read depth variation is much more complicated in bisulfite sequencing studies than in RNA-seq studies (Figure S1). Because *voom* + *limma* also cannot account for population structure, we report these results in the SI but focus on results from the simple linear model in the main text.

To compute empirical false discovery rates in simulated data, we divided the number of false positives detected at a given *p*-value threshold by the total number of sites called by the model as significant at that threshold (i.e., the sum of false positives and true positives). To compute empirical false discovery rates in the real data, in which the false positives and true positives were unknown, we used permutations. Specifically, we permuted the predictor variable for each data set four times, reran our analyses, and then calculated the false discovery rate as the average number of sites detected at a given *p*-value threshold in the permuted data divided by the total number of sites detected at that threshold in the real data. For simulated data sets only, we also calculated the area under the receiver operating characteristic curve (AUC) to produce a measure of the overall tradeoff between detecting true positives and calling false positives.

### Analysis of age-related changes in DNA methylation levels

Our initial analyses of the baboon RRBS dataset focused only on the relative ability of each method to detect age-associated sites. For these analyses, we therefore did not control for other biological covariates that may contribute to variance in DNA methylation levels (note that biological covariates cannot be incorporated into several implementations of the beta-binomial model [31,32]: see Table 1). However, to investigate patterns of age-related changes in DNA methylation levels, and to compare them to previously described patterns in the literature, we wished to control for such covariates. To do so, we reran the differential methylation analysis in MACAU, this time controlling for sex, sample age, and efficiency of the bisulfite conversion rate estimated from the lambda phage spike-in.

First, we investigated whether age-associated sites were enriched in functionally coherent regions of the genome, many of which have previously been identified as age-related [38,92,93]. To do so, we defined gene bodies as the regions between the 5’-most transcription start site (TSS) and 3’-most transcription end site (TES) of each gene using *Panu* 2.0 annotations from Ensembl [133]. We defined promoter regions as the 2 kb upstream of the TSS. CpG were annotated based on the UCSC Genome Browser track for baboon [134], with CpG island shores defined as the 2 kb regions flanking either side of the CpG island boundary (following [26,135,136]). Finally, because no enhancer annotations are available that are specific to baboons, we used H3K4me1 ChIP-seq data generated by ENCODE (from human peripheral blood mononuclear cells) to define enhancer regions [97]. In addition, we used chromatin state annotations from the Roadmap Epigenomics Project (also generated from human peripheral blood mononuclear cells) to further investigate biases in the locations of age-associated sites [96]. Using these annotation sets, we performed Fisher’s Exact Tests to ask whether age-associated sites were enriched or underrepresented in specific genomic regions. To identify differentially methylated regions (DMRs), we used the criteria proposed by [137]. Specifically, DMRs contained at least 3 differentially methylated sites with an inter-CpG distance ≤1 kb, with only 3 non-differentially methylated sites permitted in the DMR as a whole.

Second, we asked whether differentially methylated sites were more likely to fall close to blood-expressed genes. For this comparison, we drew on previously published RNA-seq data, generated from whole blood samples collected in the Amboseli baboon population [43]. We defined blood-expressed genes as those genes that had non-zero counts in more than 10% of individuals in the RNA-seq data sets, and that had mean read counts greater than or equal to 10. We then compared the number of differentially methylated CpG sites near blood-expressed genes (i.e., within the gene body or within 10 kb of the gene TSS or TES) to the number of differentially methylated CpG sites near genes that were not expressed in blood, using a Fisher’s Exact Test.

Finally, we investigated whether CpG sites that occur near genes that are differentially expressed with age were also more likely to be differentially methylated with age. For this comparison, we defined ‘age-associated genes’ as genes differentially expressed with age (at a 10% FDR) in the RNA-seq data set [43]. We compared the number of differentially methylated CpG sites near blood-expressed, age-associated genes to the number of differentially methylated CpG sites near genes that were not within this set of genes, again using a Fisher’s Exact Test.

### Ethics statement

The baboon data used in this study was generated from samples collected from wild baboons living in the Amboseli ecosystem of southern Kenya. This research is conducted under the authority of the Kenya Wildlife Service (KWS), the Kenyan governmental body that oversees wildlife (permit number NCST/RCD/12B/012/57 to Jenny Tung). As the animals are members of a wild population, KWS requires that we do not interfere with injuries to study subjects inflicted by predators, conspecifics, or through other naturally occurring events. Permission to perform temporary immobilizations (for blood sample collection) was granted by KWS; further, these immobilizations were supervised by a KWS-approved Kenyan veterinarian, who monitored anesthetized animals for hypothermia, hyperthermia, and trauma (no such events occurred during our sample collection efforts). Observational and sample collection protocols were approved though IACUC committees at Duke University (current protocol is A020–15–01 to Jenny Tung and Susan C. Alberts).

### Software and data availability

The MACAU software and a custom script for implementing a beta-binomial model in R is available at: www.xzlab.org/software.html. Previously published data sets are available at http://bergelson.uchicago.edu/regmap-data/regmap.html/ *(Arabidopsis* SNP genotype data); http://www.ncbi.nlm.nih.gov/geo/ *(Arabidopsis* WGBS data: GSE43857); http://www.nature.com/nature/journal/v465/n7298/full/nature08800.html#supplementary-information *(Arabidopsis* phenotype data); and http://www.ncbi.nlm.nih.gov/sra (Baboon RNA-seq data: GSE63788). Baboon RRBS data generated in this study are deposited in NCBI (project accession SRP058411).

## Acknowledgments

We thank the Kenya Wildlife Services, Institute of Primate Research, National Museums of Kenya, National Council for Science and Technology, members of the Amboseli-Longido pastoralist communities, Tortilis Camp, and Ker & Downey Safaris for their assistance in Kenya. We also thank Jeanne Altmann and Susan Alberts for support and access to the Amboseli data set and samples; Raphael Mututua, Serah Sayialel, Kinyua Warutere, Mercy Akinyi, Tim Wango, and Vivian Oudu for invaluable assistance with sample collection; Alexander Meissner, Joe Aman, and Joe DeYoung for assistance with RRBS; Matthew Stephens and Sayan Mukherjee for insight and support on previous versions of MACAU; and Susan Alberts, Hyun Min Kang, Sayan Mukherjee, Roger Pique-Regi, Dan Runcie, William Wen, and three anonymous reviewers for useful comments on a previous version of the manuscript. Finally, we thank the Baylor College of Medicine Human Genome Sequencing Center for access to the current version of the baboon genome assembly *(Panu 2.0*).

## Supporting Information

Text S1: Supplementary text

Figures S1-S24: Supplementary figures

Tables S1-S5: Supplementary tables

---

**Supplementary Figure 1.** In a real WGBS dataset (from *Arabidopsis)* and a real RRBS dataset (from yellow baboons), coverage varies widely across CpG sites and individuals. For each CpG site represented in each data set (n=433,871 for baboon and n=830,676 for *Arabidopsis),* we calculated the mean site-specific coverage across individuals, as well as the standard deviation of coverage values for those sites. The distribution of these values are are shown for the baboon RRBS dataset (A-B, in blue) and the *Arabidopis* WGBS dataset (C-D, in green). Average coverage values are depicted in A and C, and coverage standard deviation values are depicted in B and D.

**Supplementary Figure 2.** MACAU *p*-values are consistent across runs. QQ-plots comparing the *p*-value distributions for 3 independent runs of MACAU on the same data sets, with different simulated heritability values (Panels A, D - h^2^ = 0; Panels B, E - h^2^ = 0.3; Panels C, F - h^2^ = 0.6). Pairwise correlations between each independent run were R > 0.95 for h^2^ = 0:,R > 0.97 for h^2^ = 0.3; and R > 0.98 for h^2^ = 0.6. Distributions shown are for analyses of simulated secondary dormancy effects on DNA methylation levels in the *Arabidopsis* data set (4000 sites, n=24 accessions).

**Supplementary Figure 3.** MACAU results are robust to prior perturbation. QQ-plots comparing the results from MACAU implemented with an uninformative prior (σ^2^ ~ U(0,1), as in the main text, x-axis) versus an alternative prior (log(σ^2^) ~ U(0,1), y-axis). All analyses tested for age effects on DNA methylation levels in a simulated baboon data (based on properties of the real baboon RRBS data and age information). Sample sizes and heritabilities are shown on each plot, as are the results from a Kolmogorov-Smirnov test comparing the two distributions represented in each plot. In all cases, the simulated percent variance explained by age was set to 10%. The number of age-associated sites detected in each analysis were identical for all simulations where n=80 (10% empirical FDR), and very similar when n=50 (0.4–0.8% more age-associated sites were detected with the alternative prior than with the uninformative prior).

**Supplementary Figure 4.** The normal mixture provides an accurate approximation to the negative log gamma distribution. (A) Density plot and (B) quantile-quantile plots demonstrating that the normal mixture approximation approximates –log(Ga(r, 1)) well even in the most difficult case when r=1.

**Supplementary Figure 5.** A binomial mixed model (BMM) implemented in MACAU is more efficent than a BMM implemented in the software MCMCglmm. (A) Computation time (in hours) is plotted for datasets containing varying numbers of individuals, but each containing 100 sites. Computation time is plotted on a log10 scale in the main plot, and on a traditional scale in the inset. (B) Computation time (in hours) is plotted for a dataset containing 150 individuals, but varying numbers of sites (in thousands) as noted on the x-axis. All computation was performed on a single core of an Intel Xeon L5420 2.50 GHz processor.

**Supplementary Figure 6.** Comparisons between methods when DNA methylation levels are not heritable, and the predictor variable is binarized. To include methods that can only analyze categorical differences in DNA methylation levels between two groups, we binarized age values in our simulated RRBS datasets (individuals below median age = young versus individuals above median age = old). We compared the AUC of each method (open circles), as well as their ability to detect true positives at a 10% FDR (closed circles). For these comparisons, we used simulated datasets with a fixed h^2^ of 0 (n = 5000 sites including 500 true positives and 4500 true negatives; percent variance explained by age varies as noted in the panel headings). Results for simulations with (A) n = 50 or (B) n = 80 individuals are plotted below. Note that the right-hand y axis for the proportion of true positives detected varies depending on sample size.

**Supplementary Figure 7.** Comparisons between methods when DNA methylation levels are heritable, and the predictor variable is binarized. To include methods that can only analyze categorical differences in DNA methylation levels between two groups, we binarized age values in our simulated RRBS datasets (individuals below median age = young versus individuals above median age = old). We compared the AUC of each method (open circles), as well as their ability to detect true positives at a 10% FDR (closed circles). For these comparisons, we used simulated datasets with a fixed sample size of 80 (n = 5000 sites including 500 true positives and 4500 true negatives; percent variance explained by age varies as noted in the panel headings). Results for simulations with (A) h^2^ = 0.3 or (B) h^2^ = 0.6 are plotted below.

**Supplementary Figure 8.** MACAU outperforms linear mixed models implemented after a variety of standard data transformation approaches. We performed four different transformations on simulated baboon bisulfite sequencing count data (n = 5000 sites including 500 true positives and 4500 true negatives; percent variance explained by age = 10%; sample size = 80, h^2^ = 0.6). Below, we use QQ-plots to compare the distribution of *p*-values produced by GEMMA (operating on the transformed data) versus MACAU (analyzing the raw count data). In all panels, the observed *p*-values are plotted against quantiles for the distribution of *p*-values obtained from running each method (MACAU or GEMMA, respectively) on permuted data. We also note the proportion of simulated true positives detected by each approach (for comparison, MACAU detects 20.6% of simulated true positives in the same dataset).

**Supplementary Figure 9.** Comparison across methods when DNA methylation levels are not heritable. We compared the AUC of each method (open circles) and their ability to detect true positives at a 10% FDR (closed circles). We did so using simulated data sets (n = 5000 sites including 500 true positives and 4500 true negatives; percent variance explained by age varies as noted in the panel headings). For all simulations shown below, h^2^ was set to 0. (A) Results for simulations with n=20 individuals; (B) with n=50 individuals; and (C) with n=80 individuals. Note that the right-hand y axis for the proportion of true positives detected varies depending on sample size.

**Supplementary Figure 10.** Secondary dormancy is correlated with population structure in the *Arabidopsis* WGBS dataset. Principal components analysis on 188,093 genotyped sites with minor allele frequency >5% reveals that genetic background is correlated with secondary dormancy values. The correlation between the secondary dormancy phenotype values and the first principal component of the genetic relatedness matrix is R^2^ = 0.38, p = 7.84 × 10^−4^ (n = 24). The first principal component (PC1) explains 8.5% of the genetic variance in the data set.

**Supplementary Figure 11.** A mixed modeling approach (implemented in GEMMA) appropriately controls for genetic covariance in simulated and real WGBS data. (A, B) The distribution of *p*-values for 4000 simulated true negative sites (n = 24 accessions; effect of secondary dormancy on DNA methylation levels = 0). For each simulation, h^2^ was set to 0.3 (A) or 0.6 (B). Simulated data were analyzed with a linear model or GEMMA, and compared against the expected uniform distribution. (C) QQ-plots comparing the *p*-value distributions for (i) a model testing for effects of secondary dormancy on DNA methylation levels in real WGBS data, plotted on the y-axis; and (ii) the same model when the secondary dormancy values were permuted across individuals, plotted on the x-axis. Here, the lack of inflated test statistics in the case of the linear model is likely due to the model’s low power (see Figure S12b, for n=25). The genomic control factor, λ, is shown for each set of results.

**Supplementary Figure 12.** A mixed modeling approach (implemented in GEMMA) controls for genetic covariance in data sets that span a range of sample sizes and levels of correlation between population structure and a predictor variable of interest. Genomic control factor when simulated datasets (n=5000 sites per dataset; h^2^ = 0.6) were analyzed with either (A) a linear model or (B) a linear mixed model implemented in GEMMA. The correlation between the simulated predictor variable and the first principal component of genome-wide genotype data is plotted on the x-axis.

**Supplementary Figure 13.** MACAU controls for genetic covariance in data sets that span a range of sample sizes and levels of correlation between population structure and a predictor variable of interest. Percent of dataset associated with the predictor variable (at a 10% FDR) when simulated datasets (n=5000 sites per dataset; h^2^ = 0.6) were analyzed with either (A) a beta-binomial model or (B) a binomial mixed model implemented in MACAU. The correlation between the simulated predictor variable and the first principal component of genome-wide genotype data is plotted on the x-axis.

**Supplementary Figure 14.** Distribution of pairwise relatedness values for baboons (n=80) from a single social group, used in simulations. Approximately half of the individuals are unrelated, while a small proportion (~10%) are highly related (i.e., related at the level of half siblings or higher, r = 0.25).

**Supplementary Figure 15.** MACAU produces well-calibrated *p*-values when the simulated effect of age is set to 0. Results from 4500 simulated sites, where we set the effect of age on DNA methylation levels equal to 0 and the heritability of DNA methylation levels equal to (A) 0, (B) 0.3, or (C) 0.6. All QQ-plots compare the distribution of *p*-values produced by MACAU to the expected uniform distribution.

**Supplementary Figure 16.** MACAU provides increased power to detect age-associated sites when DNA methylation levels are heritable. We simulated age effects on DNA methylation levels, in presence of genetic effects (panel A, h^2^ = 0.3; panel B, h^2^ = 0.6) across a range of effect sizes. The proportion of true positives detected at a 10% empirical FDR is plotted for each method (closed circles) as is the AUC (open circles). For all simulations shown here, the sample size was set to 80 individuals.

**Supplementary Figure 17.** MACAU provides increased power to detect age-associated sites when DNA methylation levels are heritable. We simulated age effects on DNA methylation levels in datasets of 500 (A-B) and 1000 individuals (CD). For all simulations, we included genetic effects on DNA methylation levels (panels A and C: h^2^ = 0.3; panels B and D: h^2^ = 0.6). Below, we show the proportion of true positives detected at a 1% empirical FDR (closed circles) as well as the AUC (open circles) for each method.

**Supplementary Figure 18.** Distribution of sites covered in the baboon RRBS dataset (n = 433,871 CpG sites). (A) Absolute number of sites analyzed for a given genomic region. See *Materials and Methods* for information on how we defined each genomic region. (B) Proportion of total annotated features in the baboon genome for which a least one CpG site was analyzed in this data set.

**Supplementary Figure 19.** DNA methylation patterns in the baboon RRBS data. (A) The distributions of bisulfite conversion rates (estimated from a spike-in sample of unmethylated lambda phage DNA) and proportions of reads starting or ending with an *Msp1* digest site, for each sample. (B) Barplots showing the distribution of DNA methylation levels by genomic compartment. As expected, CpG islands, H3K3me1-marked enhancers and promoters tend to be lowly methylated, while gene bodies and the background set of all sites analyzed tend to be hypermethylated. See [1] for similar results from a human RRBS dataset. (C) For each CpG site within 5000 bp of an annotated Ensembl TSS, we calculated the mean DNA methylation level at that site across all 50 baboons. These mean levels are plotted as a smoothed function of distance from the TSS, stratified by gene expression level quartiles obtained from baboon whole blood RNA-seq [2]. As expected, more highly methylated regions are associated with more lowly expressed genes. Only expressed genes were included.

**Supplementary Figure 20.** Distribution of p-values from four different methods for the real RRBS data. QQ-plots comparing the p-value distributions for (i) a model testing for effects of age on DNA methylation levels in real RRBS data, plotted on the y-axis; and (ii) the same model when the age values were permuted across individuals, plotted on the x-axis. For each method, the number of sites detected at a 10% FDR was as follows: Beta-binomial = 747, GEMMA = 205, Linear 324, MACAU = 1018.

**Supplementary Figure 21.** MACAU detects differentially methylated regions in the baboon genome. Using the criteria of [3], we detected 142 age-related DMRs. Two representative DMRs are plotted in panels A and B (location of DMR in panel A: Chr14, 908111–908168; and panel B: Chr 20: 996106–996139; see Table S5 for the locations of additional DMRs). To detect DMRs, baboon ages were binarized into two categories, based on whether an individual’s age fell above or below the median age in our sample. Smoothed estimates of DNA methylation levels are shown for each age group, and the location of measured CpG sites are noted along the x-axis by black dots. Panel C shows the proportion of all identified DMRs that fell in a CpG island, CpG island shore, or both.

**Supplementary Figure 22.** Sites identified by MACAU are consistently enriched near genes identified as age-associated in the same population. For each method below, we asked whether CpG sites that occur near age-associated genes (identified using RNA-seq data from [2]) were more likely to be differentially methylated with age compared to the background set of all CpG sites near genes (using a Fisher’s exact test). We report the enrichment observed and show whether the p-value associated with the Fisher’s exact test (FET) was below 0.05 (triangles). We repeated this analysis using a varying number of top CpG sites from each method, with the number for each analysis shown on the x-axis. Dotted vertical lines correspond to the number of sites detected by MACAU at a 10% empirical FDR (a more conservative approach), or at a 10% FDR calculated in the R package *qvalue* [4] (a less conservative approach).

**Supplementary Figure 23.** MACAU is robust to error in the estimation of pairwise genetic relatedness. To understand how the performance of MACAU varies when there is error in the estimation of pairwise genetic relatedness, we added random error drawn from a normal distribution with mean 0 and standard deviation as shown on the x-axis. We then reran our analyses of simulated data sets with varying heritabilities (as shown in the figure legend inset) where n=80 and percent variance explained by age=10%. For each analysis, we show the number of simulated true positives detected by MACAU at a 10% empirical FDR (note that the results from our original analyses, with no error in the estimation of pairwise genetic relatedness, corresponds to the results for SD = 0 on the x-axis).

**Supplementary Figure 24.** MACAU outperforms the linear modeling approach implemented in ‘voom + limma’. We tested the performance of a commonly used method for modeling RNA-seq data: the combination of the *voom* function for data weighting with *limma,* a linear model approach [5]. We used simulated baboon bisulfite sequencing count data where the percent variance explained by age = 10%, sample size = 80, and h^2^ = 0.6 (n = 5000 sites including 500 true positives and 4500 true negatives). QQ-plots show the results for both the *voom* + *limma* approach (purple), as well as results from the same dataset using MACAU (orange). QQ-plots compare the p-value distributions for (i) a model testing for the effect of age on DNA methylation levels, plotted on the y-axis; and (ii) the same model when the age values were permuted across individuals, plotted on the x-axis (i.e., the null distribution of p-values). MACAU detects 20.6% of simulated true positives at a 10% FDR, while the *voom* + *limma* approach detects less than 1% of simulated true positives.

**Table S1.** Normal Mixture Approximations to -log(Ga(r, 1)) for r in [1, 5]. Normal mixture approximations to -log(Ga(r, 1)) for r in [1, 5]. A separate normal mixture distribution is used to approximate each negative log gamma distribution. The estimated parameters in the normal mixture distribution ensure that the Kullback-Leibler (KL) divergence between the two distributions is below 5×10^−4^. The parameters in the normal mixture distribution include the number of normal components (k), their weights (w), means (m) and variances (σ^2^). Means and variances are shown in their standardized version, where Ψ(r) denotes the diagamma function and Ψ(r) denotes the trigamma function.

**Table S2.** Normal Mixture Approximations to -log(Ga(r, 1)) for r in [6, 170]. Normal mixture approximations to -log(Ga(r, 1)) for r in [6, 170]. A separate normal mixture distribution is used to approximate each negative log gamma distribution. The estimated parameters in the normal mixture distribution ensure that the Kullback-Leibler (KL) divergence between the two distributions is below 5×10^−4^. The parameters in the normal mixture distribution include the number of normal components (k), their weights (w), means (m) and variances (σ^2^), all of which are functions of r. Means and variances are shown in their standardized version, where Ψ(r) denotes the diagamma function and Ψ(r) denotes the trigamma function.

**Table S3.** Computation times for each method on the two real datasets. Computation was performed on a single core of an Intel Xeon L5420 2.50 GHz processor. *n* = number of individuals; *m* = number of sites.

**Table S4.** Baboon RRBS dataset sample characteristics and read mapping summary.

**Table S5.** Locations of identified age-DMRs in the baboon genome.

